# MicroED structure of a protoglobin reactive carbene intermediate

**DOI:** 10.1101/2022.10.18.512604

**Authors:** Emma Danelius, Nicholas J. Porter, Johan Unge, Frances H. Arnold, Tamir Gonen

**Author notes:** **Additional information** Correspondence and requests for materials should be addressed to Tamir Gonen.

## Abstract

Microcrystal electron diffraction (MicroED) is an emerging technique which has shown great potential for describing new chemical and biological molecular structures. [1] Several important structures of small molecules, natural products and peptides have been determined using *ab initio* methods. [2] However, only a couple of novel protein structures have thus far been derived by MicroED. [3, 4] Taking advantage of recent technological advances including higher acceleration voltage and using a low-noise detector in counting mode, we have determined the first structure of an *Aeropyrum pernix* protoglobin (*Ape*Pgb) variant by MicroED using an AlphaFold2 model for phasing. The structure revealed that mutations introduced during directed evolution enhance carbene transfer activity by reorienting an alphahelix of *Ape*Pgb into a dynamic loop making the catalytic active site more readily accessible. After exposing the tiny crystals to substrate, we also trapped the reactive iron-carbenoid intermediate involved in this engineered *Ape*Pgb’s new-to-nature activity, a challenging carbene transfer from a diazirine via a putative metallo-carbene. The bound structure discloses how an enlarged active site pocket stabilizes the carbene bound to the heme iron and, presumably, the transition state for formation of this key intermediate. This work demonstrates that improved MicroED technology and the advancement in protein structure prediction now enables investigation of structures that were previously beyond reach.

## Main

The identification of novel enzymes through protein engineering and directed evolution has made biocatalysis a competitive tool in modern organic synthesis. [5] Heme enzymes are particularly interesting due to their ability to form and transfer reactive carbene and nitrene intermediates to effect transformations not known in biology, and sometimes not even known in chemical catalysis. [6-8] Although many new-to-nature heme enzymes have been described, with a wide diversity in their synthetic products, the structural rationale behind these advancements is still missing. Describing the short-lived reactive intermediates of these reactions is of great interest for development of future biocatalysts, but this has proven very challenging. Using x-ray crystallography, only two carbene-bound intermediates have been reported: a carbene-bound *Rhodothermus marinus* cytochrome c variant [9] and a myoglobin heme-iron–carbenoid complex, which was observed in a non-reactive configuration. [10] In many cases, large, well-ordered crystals of the enzymes and their complexes are not accessible, so other methods able to handle much smaller crystals are needed.

MicroED is a cryo-electron microscopy (cryo-EM) method which has been developed during the last decade and has contributed many structures ranging from small molecules [11] and peptides [12] to both soluble [1] and membrane proteins. [13] In MicroED, electron diffraction data is collected from three-dimensional crystals using a transmission electron microscope (TEM) under cryogenic conditions. The crystals are typically a billionth the size of crystals used for X-ray diffraction, hence structures of new and important targets which have been out of reach due to challenges in crystal growth can be determined. [14] As in x-ray diffraction experiments, the intensities of the diffracted beams are directly recorded while the phases also used to model the crystal content need to be derived by other means. At atomic resolution, phases can be estimated directly from the intensities computationally by *ab initio* methods. Several novel small molecules, peptides and natural products have been solved by MicroED using *ab initio* phasing, including the sub-ångström structure of the prion proto-PrPSc peptide, [15] the antibiotic macrocycle thiostrepton [11] and the chemotherapeutic teniposide. [16] Further, radiation damage-induced phasing has been shown previously for MicroED data, [17] and isomorphous replacement, while theoretically possible, has yet to be demonstrated effectively. Recently, even for macromolecular structures, *ab initio* phasing was demonstrated with the sub-ångström resolution structure of triclinic lysozyme. [18] However, like with x-ray crystallography, the most common method to derive initial phases for macromolecular MicroED structure determination is molecular replacement (MR), which relies on a starting homologous model. The model is typically a similar protein with a known structure and the phases can be calculated after its position and orientation are found within the crystal. Due to the growing number of known structures deposited to the PDB as well as computational improvements, MR usage has increased from 50% in 2000 to 80% in 2022, and as such MR is the first choice in most cases for both x-ray crystallography and macromolecular MicroED. A couple of novel protein structures have been solved by MicroED with MR using the known structure of the wild-type homolog, including a novel mutant of the murine voltage-dependent anion channel at 3.1 Å resolution, [4] and the 3.0 Å structure R2lox, [3] but MR remains challenging in cases where structures of closely related homologues are not available.

Recent advances in protein structure prediction can enable MR where experimentally determined structures fail or otherwise are unavailable. The possibility to generate *ab initio* models without closely related homologues took a leap in the 14^th^ Critical Assessment of Structure Prediction (CASP14) with the emergence of the deep learning method implemented in AlphaFold2. On the provided test sets, the peptide backbone atom positions could be predicted accurately to within 1 Å. This accuracy meets the requirement for MR when the diffracting resolution is better than 3 Å. [19] AlphaFold2 or RosettaMR have already been used to generate a starting model for successfully phasing x-ray data where no experimental structure was available. [20-22] However this approach has not been successfully applied to MicroED data.

Here, we present the previously unknown structure of *Aeropyrum pernix* protoglobin (*Ape*Pgb) determined by MicroED in two different states: resting state and with the reactive intermediate carbene bound following chemical activation of the reaction. The *Ape*Pgb structure described herein is an engineered variant for which no wild-type structure has been experimentally determined. The crystals formed as long and thin plates that were brittle and challenging to isolate; despite significant efforts, the structure could not be obtained by synchrotron x-ray crystallography as very weak or no diffraction was observed. The structure was obtained using the latest MicroED technology including a cryo-TEM operating at 300 kV acceleration with parallel illumination, data collection on a direct electron detector operating in counting mode, and cryogenic preservation. The resting state structure was solved by molecular replacement against a computationally generated model from AlphaFold2. Following exposure of the crystals to reaction-like conditions, the same methodology was used to capture and determine the structure of the carbene-bound reactive intermediate of *Ape*Pgb by MicroED. This is to the best of our knowledge, the first example of a protein structure bound to an aryl carbene intermediate. This demonstrates the feasibility of using *ab initio* generated protein models for MR in MicroED at the resolution most well-represented by protein structures in the PDB, and that MicroED can now contribute novel protein structures, including those of short-lived reactive intermediates, that were previously beyond technological reach.

### Protoglobin structural interrogation

Protoglobins are small dimeric heme proteins found in Archaea that are presumed to naturally function as gas binders/sensors. [23] These proteins have recently gained attention as engineered carbene transfer biocatalysts that can use either diazo compounds [24] or diazirines [25] as carbene precursors. Notably, the recent report of diazirine activation (Fig. 1) represents the first example of catalytic activation and subsequent carbene transfer from these species. Characterizing the structural details underlying these laboratory-evolved functions can provide deeper insights to guide the future engineering of such biocatalysts.

**Figure 1.**
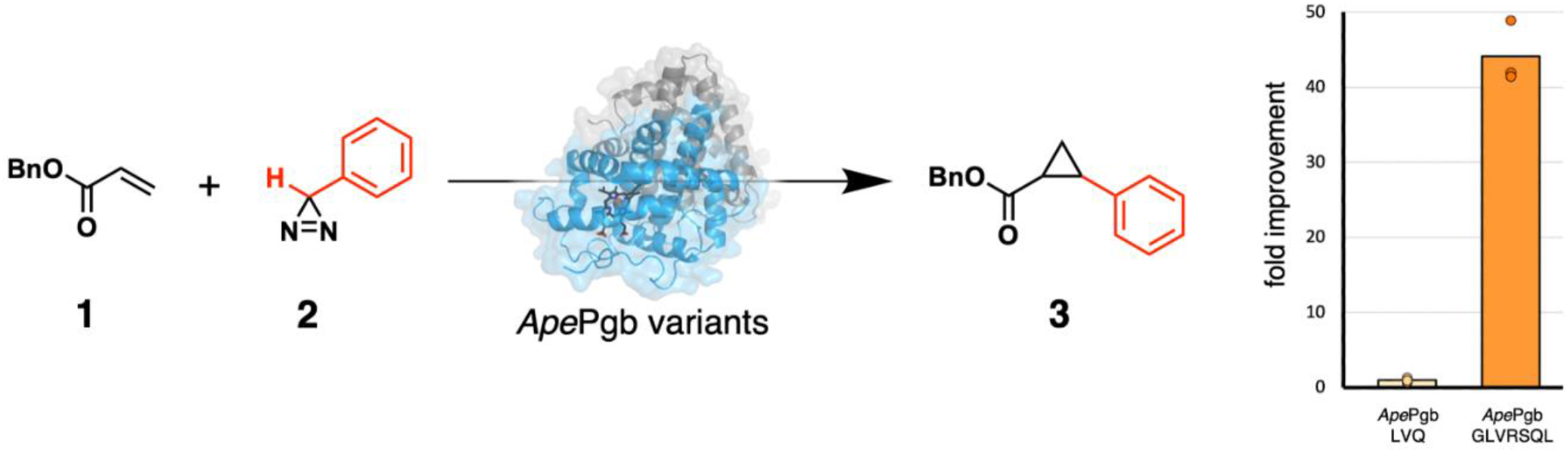
Directed evolution of *A. pernix* protoglobin (*Ape*Pgb) converted this gas-binding protein into an enzyme catalyzing cyclopropanation of benzyl acrylate **1** using phenyldiazirine **2** as a carbene source to generate cyclopropane **3**. The installation of the 4 mutations shown here, introduced during directed evolution, resulted in a >40-fold increase in activity (right; *Ape*Pgb LVQ = *Ape*Pgb W59L Y60V F145Q & *Ape*Pgb GLVRSQL = *Ape*Pgb C45G W59L Y60V V63R C102S F145Q I149L). The reaction scope was further extended to include N–H and Si–H insertion reactions. [18]

The *Ape*Pgb variant GLVRSQL described here was expressed and purified as reported previously. [25] Over 500 conditions were screened for crystal formation identifying only one condition that yielded crystals. To interrogate the structural basis for the gain of cyclopropanation activity (Fig. 1), we attempted to determine the crystal structure by x-ray diffraction (XRD). However, the crystals were extremely thin and brittle plates that formed in large clusters (Fig. 2a), making it difficult to isolate a single and intact crystal for XRD. While screening, isolated crystals diffracted weakly to around 10–12 Å resolution (Fig. 2b), proving insufficient for any structural determination. Crystal optimization assays failed to yield better crystals for XRD despite significant effort. Instead, the plate-like crystals were prepared for MicroED as described below.

**Figure 2.**
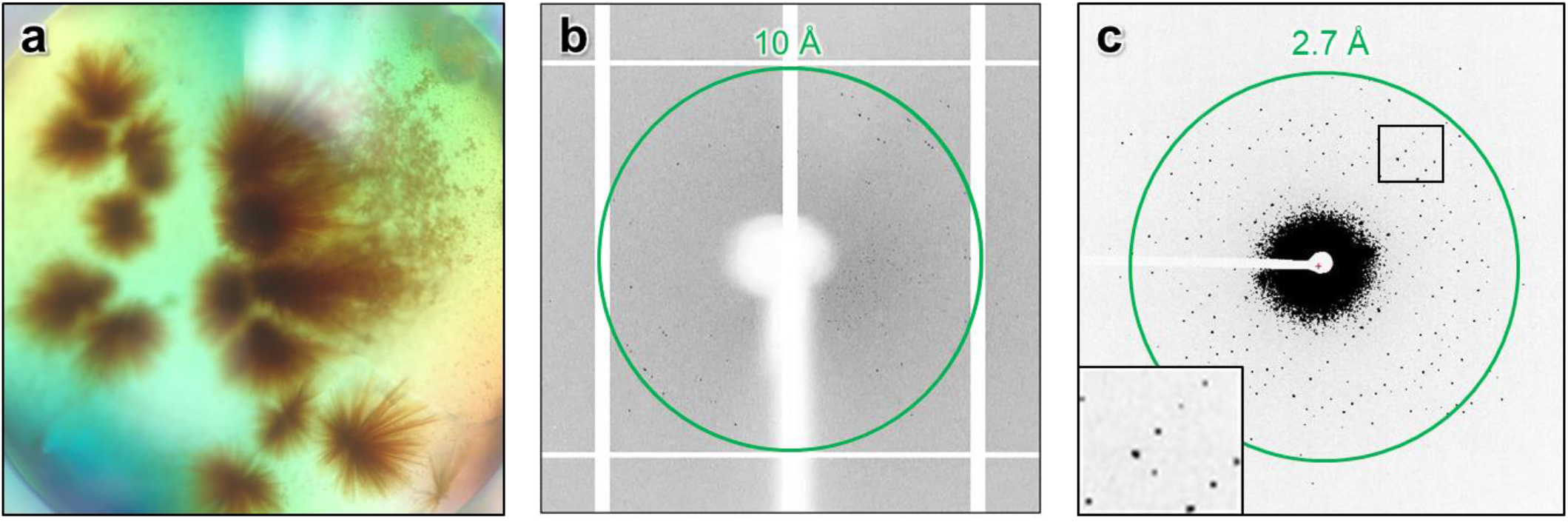
a) The crystal drop of *Ape*Pgb GLVRSQL in 0.4 M sodium phosphate monobasic / 1.6 M potassium phosphate dibasic, 0.1 imidazole (pH 8.0), 0.2 M NaCl. (b) XRD, single exposure. (c) MicroED, single exposure. Green circle indicates levels of resolution.

### MicroED grid preparation and diffraction screening

To examine the crystals in the cryo TEM, the crystalline clusters were broken into smaller crystallite fragments by perturbation using a pipette, and the remaining crystal slurry was transferred to TEM grids inside of a vitrification robot at 4ºC and 90% humidity. The grids were blotted from the back, vitrified by plunging into liquid ethane, and loaded into a Thermo Fisher Talos Arctica under cryogenic conditions for screening. The crystals appeared as thin sheets on the grids under low-magnification (Extended Data Fig. 1). Initial diffraction data were collected on a Ceta-D detector as a movie, and processed according to standard MicroED procedures. [26] However, these plate-like crystals adopted a preferred orientation on the grid and they had the low symmetry P1 space group that resulted in low completeness and the overall data quality was insufficient for structural determination.

### MicroED data collection

The data quality was dramatically improved by turning to higher acceleration voltage (300 kV) and parallel illumination at the Thermo Fisher Titan Krios TEM, and by collecting the data on the Falcon-4 direct electron detector operating in counting mode, which provides a significantly lower background and higher signal-to-noise ratio. [18] Whereas scintillator-based cameras, such as the Ceta-D used initially, record the data using integrating mode where the number of electrons is determined by the charge accumulated in a pixel during a readout-cycle, direct electron detectors such as the Falcon-4 can be used in counting mode where they detect individual electrons leading to increased accuracy and higher data quality. The higher acceleration voltage also allowed us to interrogate slightly thicker crystals, further enhancing the signal. Compared with the Ceta-D, the increased sensitivity of the Falcon-4 detector allows more information to be recorded for an identical exposure. In addition, with a faster readout more fine sliced data can be collected, further reducing the background for high resolution reflections. In this case, 840 frames were collected from each crystal on the Falcon 4, as compared with only 160 frames on the Ceta-D for the same exposure. The crystals were continuously rotated (0.15° s^−1^) in the electron beam during the exposure, covering the complete angular range of the stage. The merged data from these experiments yielded about 75% completeness with reasonable merging statistics even with P1 symmetry (Extended Data Table 1). The continuous-rotation MicroED data were converted to SMV format using an in-house software that is freely available. [27]

### MicroED data processing

Data were indexed and integrated in XDS, as described previously for MicroED. [18] The integrated data were indexed in space group P1 with unit cell dimensions (*a, b, c*) = 46.2 Å, 58.3 Å, 80.7 Å, and angles (α, β, γ) = 104.1°, 98.6°, 90.1°. Scaling and merging in AIMLESS [28] yielded a dataset to 2.1 Å resolution with an overall completeness of ∼75% at a CC_1/2_/R_merge_ of 97.2/0.18. Initially, phasing was attempted by MR using the structure of a homolog of *Ape*Pgb GLVRSQL, Y61A *Methanosarcina acetivorans* protoglobin (*Ma*Pgb Y61A; 56% identity, PDB 3ZJI). [29] However, when no reasonable solution was obtained with this starting model, we redirected our efforts towards predicted models; the sequence of *Ape*Pgb GLVRSQL was subjected to structure prediction with AlphaFold2 using the ColabFold environment. [30] The generated model was then used as a search model in Phaser to provide a preliminary phase solution for the MicroED data. The best solution with an LLG value of >1300 found 4 monomers in the unit cell. Atomic models were refined with electron scattering factors in Phenix Refine [31] using the automated solvent modeling. Several rounds of refinement resulted in a R_work_/R_free_ of 0.19/0.22. In addition to the protein with the expected heme groups, the final model includes an imidazole molecule bound to the Fe of the heme in each chain, as well as about 170 water molecules.

### Structure Analysis *Ape*Pgb GLVRSQL

The structure of *Ape*Pgb GLVRSQL (Fig. 3) has 7 mutations installed during directed evolution as compared to the wild-type sequence: C45G, W59L, Y60V, V63R, C102S, F145Q, and I149L. Most of the mutations are near the active site and affect the internal surface. The structure adopts an expanded version of the 3/3 helical sandwich typical of “classical” globins, with an additional N-terminal extension followed by the Z-helix, helping the formation of the homodimer. [29] The dimer is built by the G- and H-helices creating a four-helix bundle for the two subunits. The alignment of the AlphaFold2 model and *Ape*Pgb GLVRSQL is presented in Fig. 3a, and the alignment of the closest homolog *Ma*Pgb Y61A with the sequence alignment in Extended Data Fig. 2. The *ab initio* model from AlphaFold2 resulted in slightly smaller overall differences to *Ape*Pgb (r.m.s.d. 0.50–0.55 Å versus 0.65–0.7 Å for *Ma*Pgb Y61A). When compared to the standard protoglobin fold observed in *Ma*Pgb Y61A, the major difference is the disruption of the B helix between residues 60–70. These residues adopt a rigid helical conformation in AlphaFold2 and *Ma*Pgb, but are found to be restructured as a loop in *Ape*Pgb GLVRSQL (Fig. 3b). Given that the Y61A mutation in *Ma*Pgb does not alter the helical conformation of this region and there is no substantial deviation in the wild-type sequences before the B helix terminus, it is reasonable that this helix would still be present in wild-type *Ape*Pgb. For the *Ape*Pgb GLVRSQL structure with four chains (A–D), the backbone of the disrupted B helix 60–70 is observed for chains A–C, but the density is substantially weaker than surrounding regions. In chain D, residues of the disrupted B helix 60–70 could not be modeled. The weaker electrostatic potential map, the slight differences between chains, and the *B*-factors of the loop (Extended Data Fig. 3) suggest that this region is flexible and thus might be able to adopt different conformations in solution. It is likely that the observed structural change stems from the mutation V63R which resulted in a 14-fold boost in product yield for the cyclopropanation reaction (Fig. 1), the largest improvement from any single mutation during enzyme engineering. In *Ma*Pgb, V63 is pointing towards the active site, where the natural substrate is two atoms only. The inclusion of the much bulkier arginine residue in this position is difficult to model within the structure of *Ape*Pgb GLVRSQL without major rearrangements. The AlphaFold2-predicted model of *Ape*Pgb GLVRSQL incorrectly orients R63 into the enzyme active site (Fig. 3b), similar to the *Ma*Pgb structure. The limited space at the active site together with repulsion effect between the positively charged iron and arginine could be the reasons for breaking up the helical conformation in this region to produce a conformation where R63 is instead positioned at the surface pointing outwards (Fig. 3b). Thus, the effects of unfavorable steric and electrostatic interactions between the heme and positively-charged arginine side chain are presumed to drive the rearrangement of residues 60–70, truncating the B helix in this variant and expanding the active site cavity (Figs. 3 c and d).

**Figure 3.**
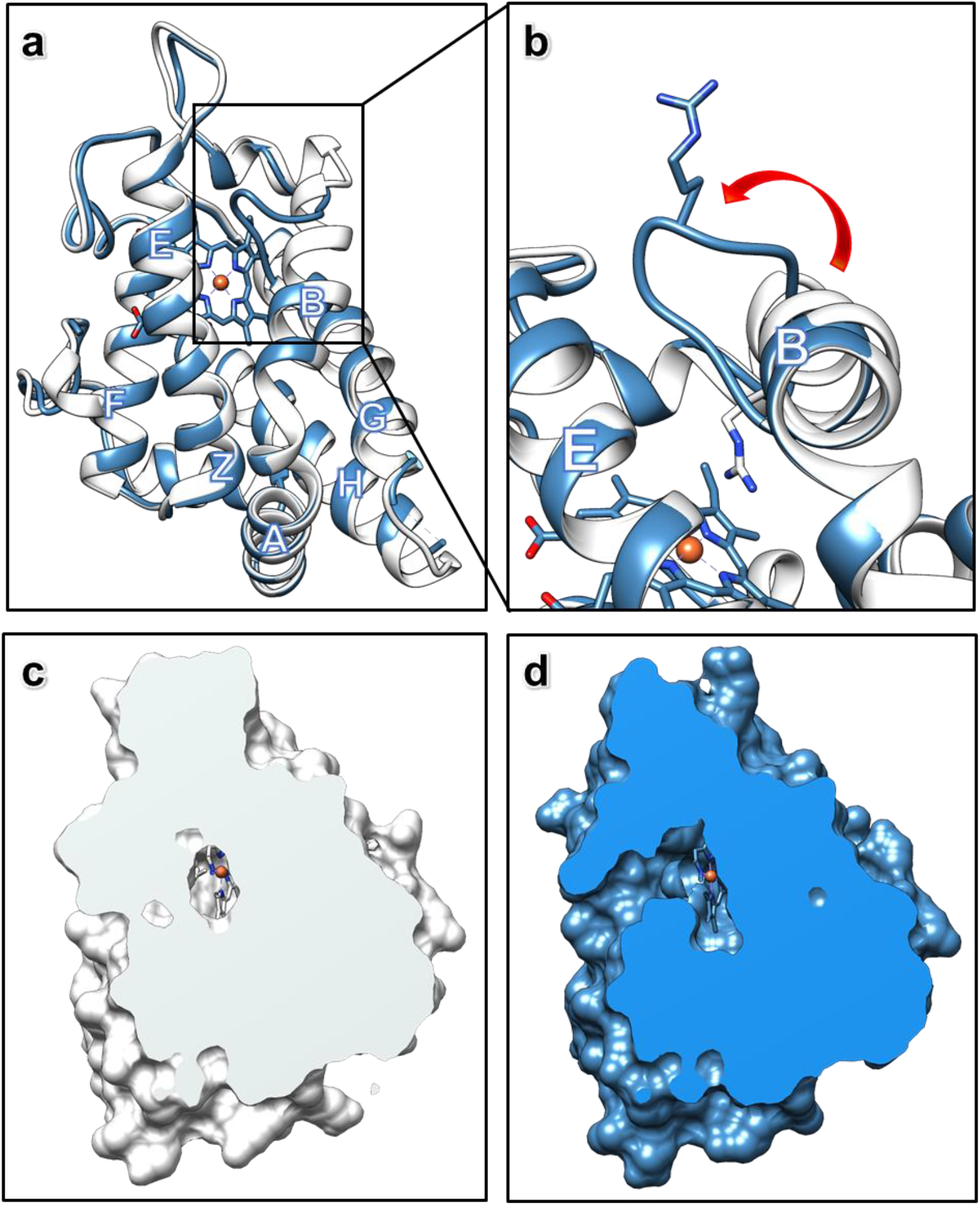
MicroED structure of *Ape*Pgb GLVRSQL: a) Superposition of the structures of *Ape*Pgb GLVRSQL (blue) with the AlphaFold2 model (white), with helices labeled according to convention for the protoglobin fold. b) Close-up of the structures of *Ape*Pgb GLVRSQL (blue) with the AlphaFold2 model (white) showing the unwinding of the B helix into a dynamic loop, creating a larger cavity around the active site with increased access to the heme. c) and d) Clipped surface of the AlphaFold2 model (white) and *Ape*Pgb GLVRSQL (blue) showing the effects of the unwinding of the helix and the rearrangement of the B/G interface leading to easier substrate access from outside in addition to the increased space available at the heme.

The solvent-inaccessible heme is buried in the protein matrix (Figs. 3c and d). This feature is in contrast to most members of the globin family structures where about 30% of the heme would be surface accessible. [23] In natural protoglobins, the diatomic substrates access the active site through two small orthogonal apolar tunnels defined at the interfaces of the B/E and B/G helices. In *Ape*Pgb GLVRSQL, however, the rearrangement of the final turns of helix B obstructs the B/E tunnel through interactions with the main chain and the W62 side chain, resulting in a broadening of the B/G tunnel (Fig. 3d). This larger tunnel is presumed to increase diffusion in the active site and allow the entry of larger ligands than the natural diatomic substrates, such as the diazirine and acrylate substrates targeted in directed evolution (Fig. 1). In fact, the bulky benzene moiety is too large to fit into the tunnels present in the AlphaFold2 model, and it seems likely that the drastic expansion of the access pathway is necessary for passage of the substrate. Molecular dynamics simulations suggest that F145 controls the accessibility of B/G tunnel, though this has yet to be validated experimentally. [23] The corresponding mutated amino acid Q145, as well as L149, in the engineered *Ape*Pgb line the expanded tunnel and could reasonably affect the affinity or orientation of the substrate. The F145Q mutation remained throughout directed evolution of the enzyme, despite screening of mutations at this position, and I149L doubled the biosynthetic yield of cyclopropane **3**, underscoring the importance of these mutations to the new activity. The mutations G45 and S102 are both located at the surface of the protein. These mutations remove the cysteine residues in the wildtype sequence. Since these cysteines are located close to one another in space in *Ma*Pgb (5.6 Å Cα–Cα separation), they are potentially capable of forming a disulfide between the A and E helices in wild type *Ape*Pgb.

### Data collection of substrate-bound *Ape*Pgb GLVRSQL

Nanocrystals allow efficient and homogeneous diffusion of small molecules, giving a fast and convenient way for the determination of ligand-bound complexes, in contrast to a time-consuming and often inaccessible co-crystallization approach. This is especially essential for ligands or intermediates with a limited half-life in solution. Hence, MicroED shows potential for structural determination of reactive intermediates in enzyme catalyzed reactions. To investigate the reactive intermediate of the reaction shown in Fig. 1, the crystal fragments were soaked with carbene precursor **2** (phenyldiazirine, Fig. 1) according to previously described protocols. [9] Following 15 minutes of soaking and the addition of sodium dithionite, the grids were prepared and data were collected as described above. The integrated data in this case were indexed in space group P121 with unit cell dimensions (a, b, c) = 58.15 Å, 45.89 Å, 71.71 Å, and angles (α, β, γ) = 90.00°, 105.42°, 90.00°. Scaling and merging in AIMLESS [28] to 2.5 Å resolution gave a dataset with overall completeness of 72% and an CC_1/2_/R_merge_ of 0.97/0.23. The data were phased by molecular replacement in Phaser using chain A of the *Ape*Pgb GLVRSQL described here. The solution found was top ranked with an LLG of 4002, containing 2 monomers in the asymmetric unit. The structure was further refined in Phenix Refine [31] to an R_work_/R_free_ of 0.23/0.28. The final model was derived by altering the angle and distance describing the carbene interaction until the lowest R_free_ value was obtained.

### Structure analysis of the metallo-carbene structure

Carbene transfer from a diazirine is thought to involve the formation of a putative reactive iron heme-carbene intermediate which transfers the carbene to a second substrate, followed by product release and regeneration of the catalyst. [25] In the metallo-carbene structure described here, the observed overall fold for the carbene-bound *Ape*Pgb GLVRSQL is the same as for the unbound, with the only small differences observed in the 60–70 loop region (Fig. 4a). Interestingly, the MicroED density of this loop is a little more defined than in unbound *Ape*Pgb, which might indicate that the loop rigidifies upon substrate binding. In particular, residue W62 is better described by the density. *Ape*Pgb GLVRSQL was engineered for activation of a benzene-substituted diazirine (Fig. 1), a much larger molecule than any natural protoglobin substrate. As discussed above, increased diffusion into and out of the active site and accommodation of larger substrates near the heme cofactor likely play a significant role in the improved activity of *Ape*Pgb GLVRSQL. For example, amino acids L59 and V60 are both located in the active site with their side chains pointing towards the binding area on the distal side of the heme group (Fig. 4b). Both the selected mutations W59L and Y60V introduce substantially smaller side chains, forming a larger cavity between the heme and the B helix. The main chain conformations for residues 59 and 60 in *Ape*Pgb GLVRSQL and the AlphaFold2 model are similar, and modeling of the original residues, W59 and Y60, suggests significant steric clashes of such residues with the aryl ring of the carbene. Further, the side chain of F93 adjusts slightly to form a pi-stacking interaction with the phenyl group of the carbene (Fig. 4b). These intermolecular interactions likely stabilize the binding and orientation of the phenyl carbene, each contributing to the improved reactivity gained through evolution. The observed MicroED density and the occupancy of the carbene suggests a single carbene species as the dominant form, where the rate of carbene formation is greater than the rate of carbene decay in absence of the second substrate.

**Figure 4.**
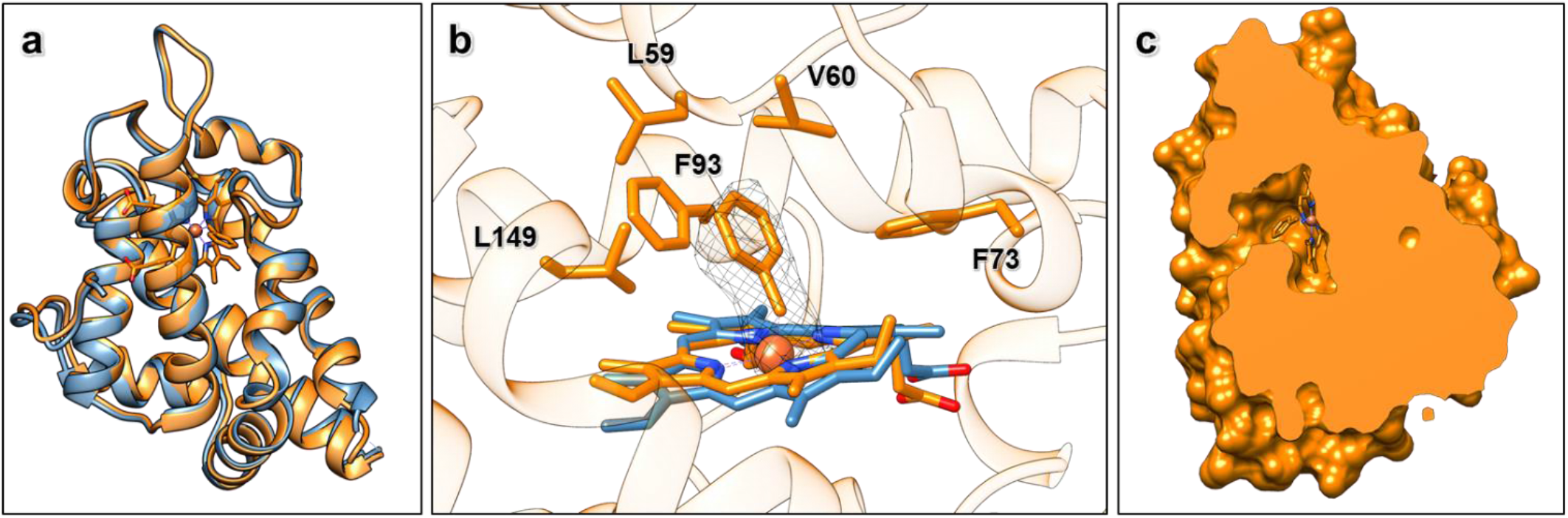
MicroED structure of carbene-bound *Ape*Pgb GLVRSQL: a) Superposition of the structures of *Ape*Pgb GLVRSQL (blue) the carbene-bound intermediate (orange). b) Polder omit map (2.5σ) for the metallo-carbene complex (orange), also including the heme from unbound *Ape*Pgb GLVRSQL (blue) for comparison. c) Clipped surface of the carbene-bound *Ape*Pgb GLVRSQL, showing a decrease in the size of the B/G channel as compared to the unbound *Ape*Pgb GLVRSQL.

While it is clear that mutations in *Ape*Pgb GLVRSQL have reshaped the active site, the heme exhibits a similar ruffled distortion to that observed in *Ma*Pgb and other protoglobins (Extended Data Fig. 2). [23] Out-of-plane distortions to the porphyrin ring are known to alter the electrostatic and ligand-binding properties of the bound iron, [32] but the specific changes associated with any specific distortion are challenging to measure and remain unclear. When comparing the *Ape*Pgb GLVRSQL with the carbene-bound intermediate, the ruffling of the heme is changed by up to 0.5 Å (Fig. 4b). The position of the carbene that resulted in the lowest R_free_ in similar refinement rounds is found at a distance of 1.74 Å (Fe – C1) and at an angle of about 128 degrees (Fe – C1 – C2). These values are comparable to the previously determined protein structure describing a heme–carbene complex, [9] as well as an iron porphyrin X-ray structure in which a diaryl-carbene is bound to the Fe atom. [33] When comparing the B/G helix interfaces of the unbound and carbene bound states, it seems that binding the substrate has closed the passage slightly (Fig. 4c), which coincides with the observation that the residues around the active site and at the solvent tunnel are less dynamic when the substrate is bound. This change is also observable in the *B*-factor gradient (Extended Figure 3). The efficiency of enzymes in accelerating chemical reactions is explained by both their ability to pre-organize the active site for transition state stabilization [34] as well as sample the conformational ensemble required for substrate binding, reaction, and product release. [35] For this, some inherent flexibility of the enzyme structure is required. The increase in flexibility observed for *Ape*Pgb GLVRSQL can enable the enzyme to adopt the conformations important for the different processes. The following observed rigidification upon binding the substrate might function to pre-organize the active site for transition state stabilization. Notably, donor-substituted carbenes are known to be short-lived and highly reactive. [36] That such a sensitive intermediate can be trapped and observed by MicroED underscores the value of this technique and the insights it can provide into such systems. The homogeneity of the bound intermediate within the crystal is likely enhanced by the improved diffusion and smaller sample size inherent to microcrystalline samples, providing better context for the atomic details underlying the enzyme chemistry. The atomic details underlying the engineered carbene transfer chemistry developed in these protoglobins will serve to guide future enzyme engineering, leading to further development of future biocatalysts.

In conclusion, comparisons to both the experimental structure of the related *Ma*Pgb as well as the predicted AlphaFold2 model show good overall agreement. It highlights the significance of the disruption introduced into the B helix region of the protoglobin fold and implicate the V63R mutation as a factor in this structural change. The broadening of the active site access tunnel relates well to the increased reaction rates observed for this variant. In modern crystallography, most protein structures are phased by molecular replacement using a related model from the Protein Structure Databank. To date, structure determination using MicroED in the absence of a reasonable search model has been set back due to the lack of experimental phasing techniques analogous to anomalous scattering in X-ray crystallography. We present the determination of a novel structure that could be solved by molecular replacement made possible by an *ab initio* generated model from AlphaFold2 in concert with higher quality data accessible due to advanced detector development and a high voltage electron microscope. We further used this technology to investigate the formation of the reactive metallo-carbene and describe the first structure of an aryl-carbene intermediate in a protein structure. As the crystals used in this study were not amenable for X-ray diffraction, this example adds an important tool for the determination of highly-sought protein structures.

## Methods

### Protein crystallization

The crystal drops used for this study were setup using 0.5 µL of 20 mg/mL protein (50 mM potassium phosphate (pH 8.0), 150 mM NaCl) and 2.5 µL precipitant (0.4 M sodium phosphate monobasic / 1.6 M potassium phosphate dibasic, 0.1 M imidazole (pH 8.0), 0.2 M NaCl). Large crystal clusters appeared after 1-3 days.

### Grid preparation, ApePgb GLVRSQL

Quantifoil R 2/2 - 200 copper mesh grids were glow discharged (30 s, 15 mA, negative polarity) and transferred to a Leica GP2 vitrification robot with the sample chamber set to 90% relative humidity and 4 °C. The crystal drop was diluted with 15 µL of mother liquor and the needle-like crystals were broken into smaller pieces by gently pipetting into the drop. The resulting protein crystal slurry (2 µL) was pipetted onto the carbon side of the grid in the vitrification chamber and allowed to incubate for 20 seconds. The grid was blotted from the back for 30 seconds, plunged into liquid ethane and transferred to liquid nitrogen for storage.

### Grid preparation, carbene-bound ApePgb GLVRSQL

The crystal drop was diluted with 10 µL 3-phenyl-3H- diazirine dissolved in the crystallization condition, making a final concentration of 30 µM 3-phenyl-3H-diazirine. The needle-like crystals were broken into smaller pieces by gently pipetting into the drop, and the resulting slurry was left at room temperature for 15 minutes. Quantifoil R 2/2 - 200 copper mesh grids were glow discharged (30 s, 15 mA, negative polarity) and transferred to a Leica GP2 vitrification robot with the sample chamber set to 90% relative humidity and 4 °C. Following addition of 50 mM sodium dithionite to the drop, 2 µL of the protein crystal slurry soaked with 3-phenyl-3H-diazirine was pipetted onto the carbon side of the grid in the vitrification chamber and allowed to incubate for 20 seconds. The grid was blotted from the back for 30 seconds, plunged into liquid ethane and transferred to liquid nitrogen for storage.

### Data collection

The grids were loaded into a Thermo Fisher Scientific Titan Krios G3i transmission electron microscope (300 kV) under cryogenic conditions. Following screening for crystals at low-magnification in imaging mode, using the Thermo Scientific EPU-D software, identified crystals which appeared as thin rectangular sheets on the grid were tested for initial diffraction using 1 second single exposure in diffraction mode. Well diffracting crystals were setup for data collection; the crystals were brought to eucentric height, and a single 1 second exposure was repeated at the starting tilt angle. The MicroED data were collected using a Falcon 4 direct electron detector in counting mode as a movie with continuous rotation of the stage at a rate of 0.15° s-1, with the selected area aperture of 100 µm and the beam stop inserted. Frames were read out every 0.5 s giving MRC datasets of 840 images, corresponding to a 60° wedge from each crystal. The total wedge that was collected over several datasets corresponded to approximately +70° to -70°.

### Data processing and refinement

The MRC files were converted to individual frames in SMV format using the freely available MicroED software (https://cryoem.ucla.edu/). The reflections were indexed and integrated in XDS. The generated datasets were scaled in AIMLESS and phased by molecular replacement in Phaser. For *Ape*Pgb GLVRSQL a structure of predicted by AlphaFold2 through the ColabFold environment was used as a search model, and for the carbene-bound structure Chain A of *Ape*Pgb GLVRSQL was used as search model. All models were refined in phenix.refine using electron-scattering factors. The statistics are given in Extended Data Tables 1 and 2.

## Acknowledgements

E.D. thanks The Wenner-Gren Foundations for their support through the Wenner-Gren Postdoctoral Fellowship. This study was supported by the National Institutes of Health P41GM136508. The Gonen laboratory is supported by funds from the Howard Hughes Medical Institute.

N.J.P. thanks Merck and the Helen Hay Whitney Foundation for their support through the Merck-Helen Hay Whitney Foundation Postdoctoral Scholarship. This publication is based on work supported by the United States Army Research Office under Contract W911NF-19-0026 for the Institute for Collaborative Biotechnologies and the G. Harold and Leila Y. Mathers Charitable Foundation.

## Author contributions

N.P. conducted protein expression and crystallization experiments. E.D. prepared the samples and conducted MicroED data collection. E.D. and J.U. analyzed the data and solved the structures. E.D., J.U., N.P, F.A, and T.G. took part in preparation of the manuscript.

## Extended Data tables

**Extended Data Table 1.** MicroED Data collection and refinement statistics, *Ape*Pgb GLVRSQL.

| Integration |  |
| --- | --- |
| Wavelength (Å) | 0.0197 |
| Acceleration voltage | 300 kV |
| Temperature | -196°C |
| Space group | P1 |
| Unit cell (a, b, c) (Å) | 46.2, 58.3, 80.7 |
| Unit cell ($\alpha = \beta = \gamma$) (°) | 104.1, 98.6, 90.1 |
| R<sub>merge</sub> | 0.18 |
| I/$\sigma$(I) | 6.96 (2.4) |
| Completeness (%) | 74 |
| CC<sub>1/2</sub> | 97.2 |
| Wilson *B*-factor (Å<sup>2</sup>) | 8.58 |
| Refinement |  |
| Resolution range (Å) | 24.47 - 2.10 |
| Total reflections (no.) | 33825 |
| R<sub>work</sub> | 0.188 |
| R<sub>free</sub> | 0.224 |
| No. of atoms | 6547 |
| Protein | 6160 |
| Ligand/ion | 192 |
| Water | 195 |
| *B*-factors |  |
| Protein | 27.25 |
| Ligand/ion | 19.81 |
| Water | 17.74 |
| R.m.s. deviation |  |
| Bonds length (Å) | 0.013 |
| Bond angles (°) | 1.464 |

**Extended Data Table 2.** MicroED Data collection and refinement statistics, *Ape*Pgb GLVRSQL metallo-carbene complex.

| Integration |  |
| --- | --- |
| Wavelength (Å) | 0.0197 |
| Acceleration voltage | 300 kV |
| Temperature | -196°C |
| Space group | P121 |
| Unit cell (a, b, c) (Å) | 58.15, 45.89, 71.71 |
| Unit cell (α, β, γ) (°) | 90.00, 105.42, 90.00 |
| R<sub>merge</sub> | 0.18 |
| I/σ(I) | 5.73 (2.6) |
| Completeness (%) | 70 |
| CC1/2 | 96.2 |
| Wilson B-factor (Å<sup>2</sup>) | 6.8 |
| Refinement |  |
| Resolution range (Å) | 24.36 - 2.50 |
| Total reflections (no.) | 40105 |
| R<sub>work</sub> | 0.235 |
| R<sub>free</sub> | 0.284 |
| No. of atoms | 3228 |
| Protein | 3104 |
| Ligand/ion | 100 |
| Water | 64 |
| B-factors |  |
| Protein | 20.81 |
| Ligand/ion | 16.43 |
| Water | 16.34 |
| R.m.s. deviation |  |
| Bonds length (Å) | 0.005 |
| Bond angles (°) | 0.877 |

## Extended Data Figures

**Extended Figure 1.**
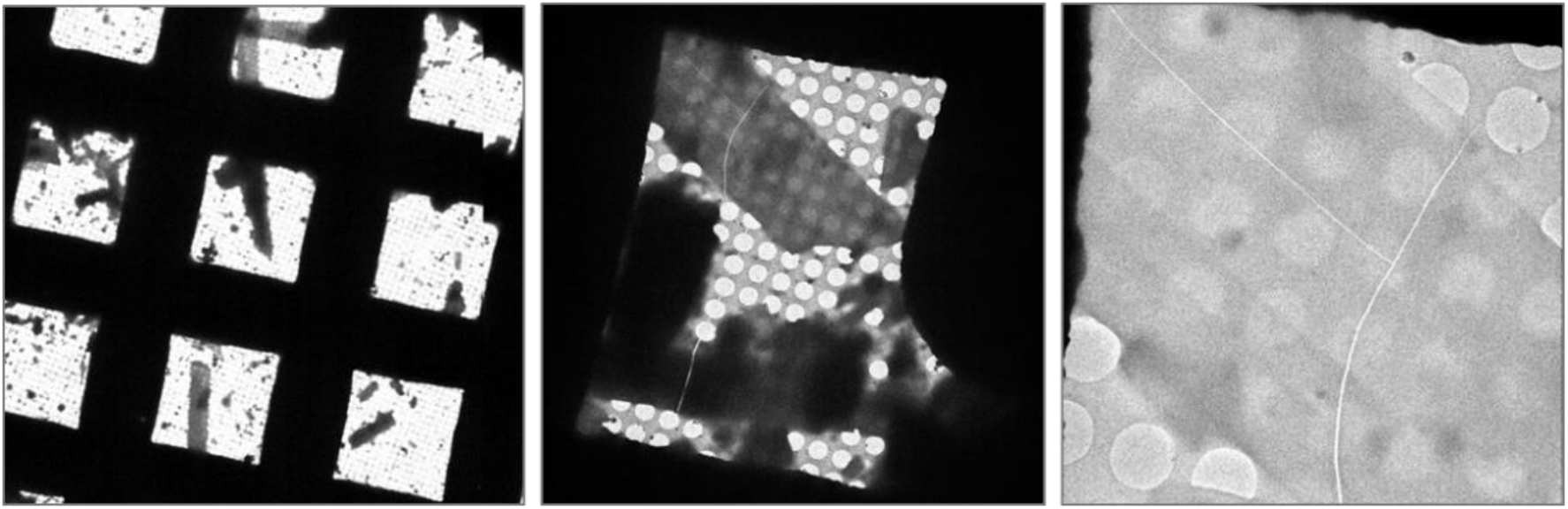
The thin plate clusters of *Ape*Pgb GLVRSQL, in the TEM at 210x, 940x, and 3400x magnification.

**Extended Figure 2.**
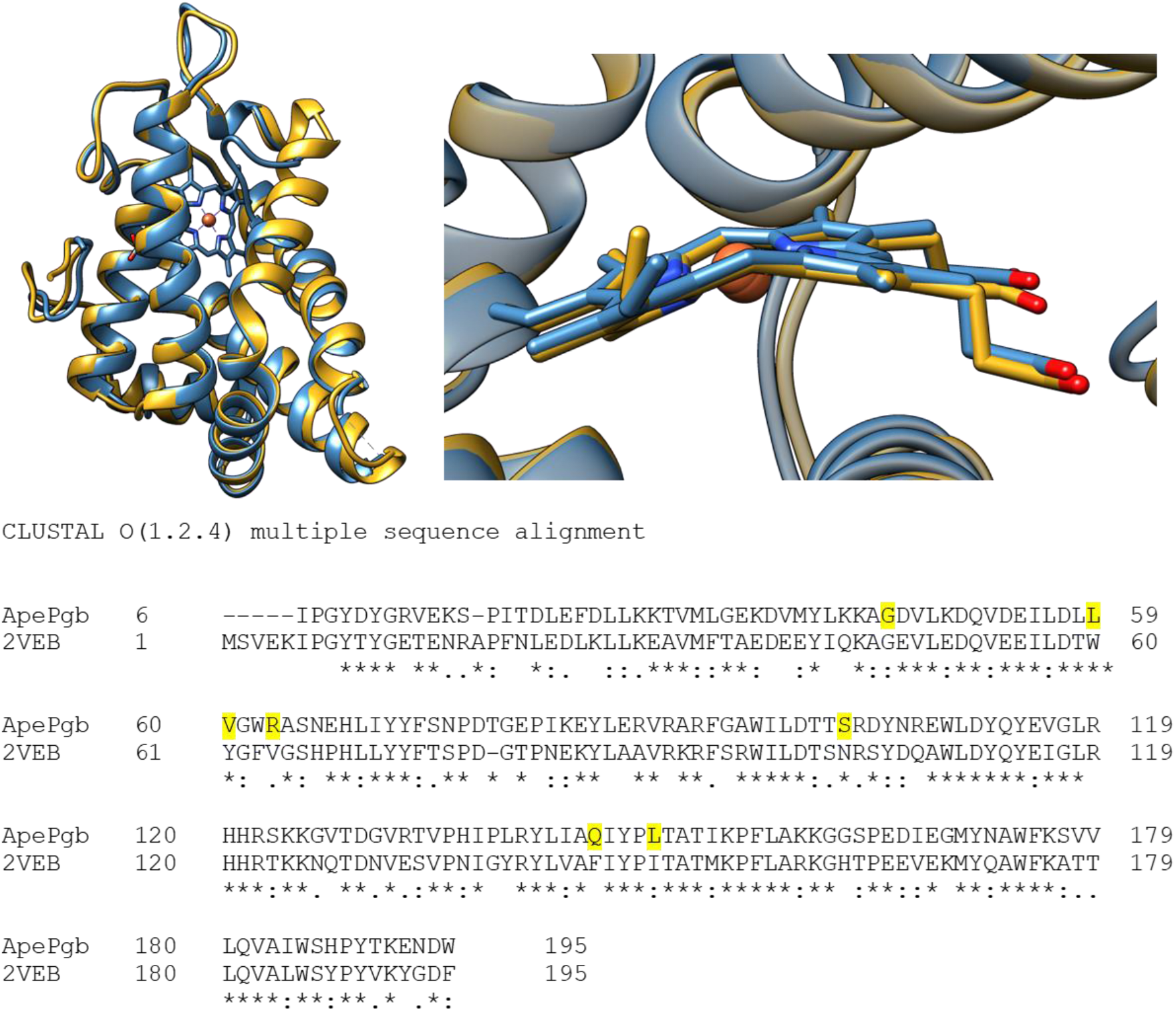
Sequence alignment and superposition of *Ape*Pgb GLVRSQL (blue) and the closest homologue protoglobin (Y61A *Methanosarcina acetivorans* protoglobin *(Ma*Pgb), PDB 3ZJI, yellow). The mutations introduced through directed evolution are highlighted in yellow. The close-up of the heme shows the similar ruffled distortion observed in *Ma*Pgb and other protoglobins.

**Extended Figure 3.**
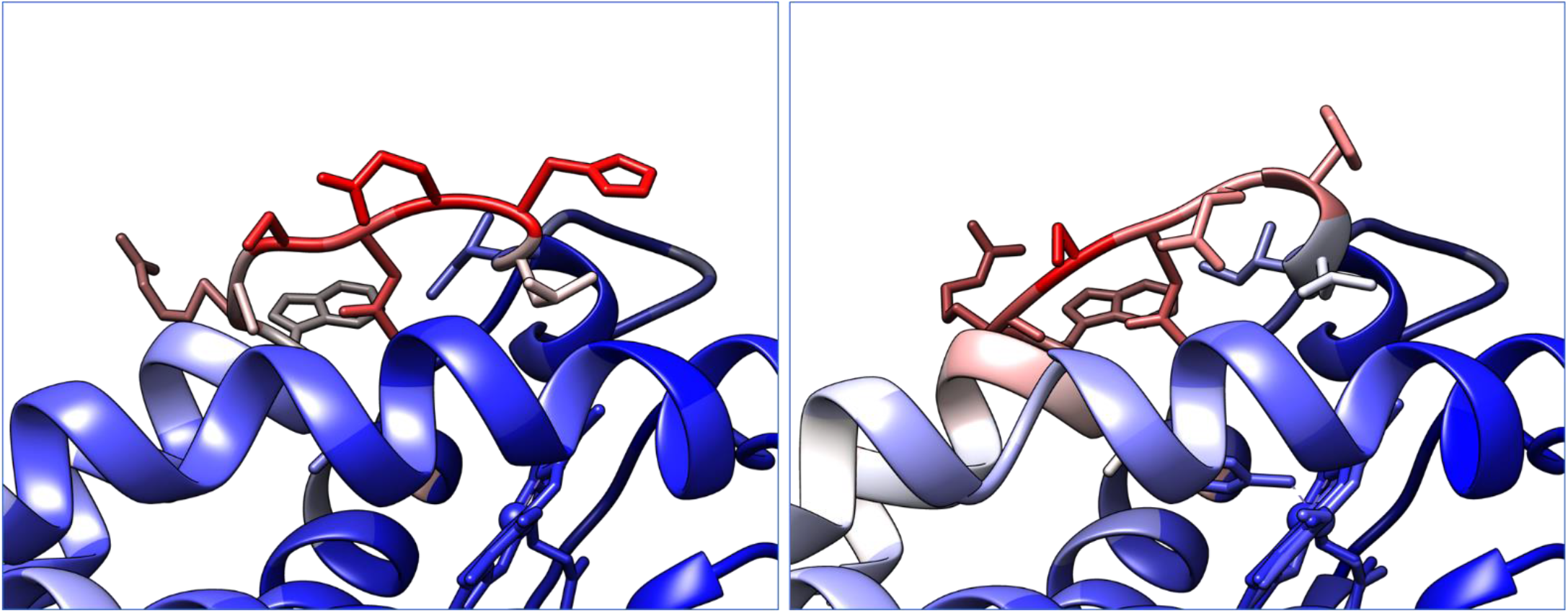
*Ape*Pgb GLVRSQL (left) and carbene-bound *Ape*Pgb GLVRSQL (right) showing the average *B*-factors as a color gradient from blue (*B*-factor 20 or less) to red (*B*-factors 50 or more) in order to highlight the *B*-factor variations within the protein.

